# Mutual dependence of Osbp and PI4KII in the maturation of regulated secretory granules

**DOI:** 10.1101/2023.07.30.551178

**Authors:** Cheng-I Jonathan Ma, Julie A. Brill

**Affiliations:** Cell Biology Program, The Hospital for Sick Children, PGCRL Building, Room 15.9716, 686 Bay Street, Toronto, Ontario, M5G 0A4, Canada; Institute of Medical Science, University of Toronto, Medical Sciences Building, Room 2374, 1 King’s College Circle, Toronto, Ontario M5S 1A8, Canada; Department of Molecular Genetics, University of Toronto, Medical Sciences Building, Room 4396, 1 King’s College Circle, Toronto, Ontario, M5S 1A8, Canada

**Keywords:** secretory granule, organelle biogenesis, autophagy, phosphatidylinositol 4-phosphate, cholesterol

## Abstract

Secretory granules (SGs) are crucial for normal animal physiology due to their role in regulated exocytosis of biologically active molecules. SG membranes are enriched in phosphatidylinositol 4-phosphate (PI4P) and cholesterol, and previous studies suggest lipid composition is important for SG biogenesis and function. Nonetheless, the molecular details of how lipids are regulated during SG biogenesis remain poorly understood. Here, we identify Oxysterol binding protein (Osbp) as a novel regulator of SG biogenesis in a *Drosophila* model. We show Osbp expression level positively correlates with SG size and that Osbp requires type II phosphatidylinositol 4-kinase (PI4KII) to increase SG size. Moreover, Osbp is needed for proper PI4KII and PI4P distribution, autophagic resolution and formation of cholesterol-rich endosomal tubules that are positive for PI4KII. Feeding larvae food supplemented with sterol leads to partial suppression of SG size and PI4P distribution defects in *Osbp* mutants. Our results indicate that reciprocal regulation of Osbp and PI4KII drives formation of membrane tubules that mediate SG maturation through elevating PI4P levels on SG membranes.

**Graphical Abstract:** 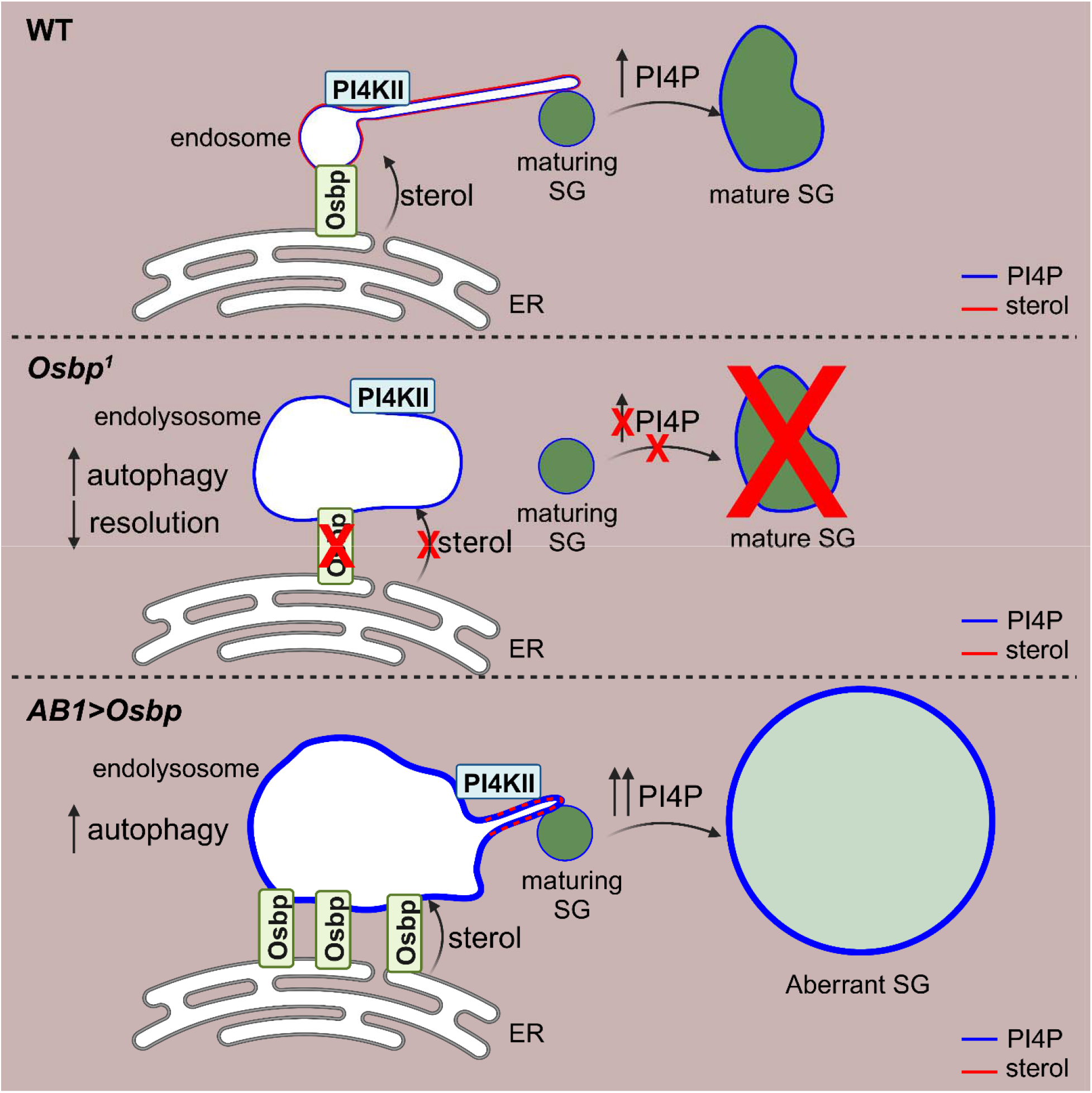

**Highlights:**

- Osbp is needed for formation of PI4KII-positive endosomal tubules that enrich secretory granule membranes with PI4P and facilitate granule maturation.
- Reciprocally, PI4KII is needed for Osbp-mediated secretory granule growth in salivary gland cells.
- Changes in Osbp expression levels alter autophagy initiation and autophagosome resolution in salivary gland cells.
- PI4KII-positive endosomal tubules are enriched in sterols, and sterol feeding suppresses defects caused by loss of Osbp.

## Introduction

The secretory pathway is essential in all eukaryotic cells because it allows for maintenance of plasma membrane integrity, delivery of integral membrane proteins and release of secretory molecules. During secretion, cargo passes through the Golgi before being trafficked to the plasma membrane via constitutive or regulated secretory pathways. Unlike constitutive secretion, cargoes in the regulated secretory pathway are stored in long-lasting SGs that require a stimulus for their release. *Drosophila* larval salivary glands provide an excellent genetic model for studying SG biogenesis in the regulated secretory pathway (Ma et al., 2021). *Drosophila* larvae stop feeding 24 h after they enter the third instar larval stage, and concurrently the salivary glands start producing glue proteins that are packaged into SGs. These SGs mature over the next 18 h before they undergo synchronized exocytosis in response to a pulse of the steroid hormone ecdysone (Biyasheva et al., 2001; Burgess et al., 2011; Kamalesh et al., 2021; Nagy et al., 2022; Rousso et al., 2016; Tran et al., 2015). Because SGs store glue proteins, their release allows the pupal case to adhere onto a solid surface during metamorphosis. Mature SGs in *Drosophila* salivary glands are 2- to 4-fold larger in cross-sectional area than immature SGs, enabling the use of fluorescently-tagged cargo proteins (Sgs3-DsRed or Sgs3-GFP) to visually identify genes required for SG biogenesis and maturation (Boda et al., 2023; Burgess et al., 2011, 2012; Ji et al., 2018; Ma and Brill, 2021a; Ma et al., 2020; Neuman et al., 2021; Reynolds et al., 2019; Syed et al., 2022; Torres et al., 2014).

Using this system, we previously showed *Drosophila* PI4KII, a type II PI 4-kinase that produces PI4P, is needed for normal SG maturation in larval salivary glands (Burgess et al., 2012). We also determined that Past1 (*Drosophila* ortholog of EHD1) and Syx16 act in the same pathway as PI4KII and are needed for formation of stable endosomal tubules positive for PI4KII and PI4P that facilitate accumulation of PI4P on SG membranes (Ma et al., 2020). The mammalian PI4KII ortholog PI4KIIα is palmitoylated and requires membrane cholesterol for its localization (Barylko et al., 2009; Lu et al., 2012). Furthermore, membrane cholesterol levels are positively correlated with PI4KIIα enzymatic activity (Minogue et al., 2010; Waugh et al., 2006). Therefore, we hypothesized that Osbp, a PI4P-binding protein involved in non-vesicular transport of cholesterol (Im et al., 2005; Mesmin et al., 2013; Raychaudhuri et al., 2006), could be important for PI4KII-mediated SG maturation.

## Results

To investigate the importance of Osbp in SG maturation, we examined an *Osbp* null mutant (*Osbp^1^*) (Ma et al., 2010, 2012). Using the SG marker Sgs3-GFP, we discovered that *Osbp^1^* mutants exhibited a small granule phenotype similar to but milder than *PI4KII* null mutants (*PI4KII*Δ; Figures 1A-1D, S1A, and S1B). Interestingly, we observed a subpopulation of SGs that became abnormally large when we overexpressed Osbp-YFP in wild-type (WT) larval salivary glands, suggesting Osbp activity could increase SG size (Figures 1E, 1F, and 1H). To test whether PI4KII activity is needed for Osbp to stimulate SG maturation, we overexpressed Osbp-YFP in *PI4KII*Δ salivary gland cells and observed that Osbp-YFP failed to increase SG size (Figures 1G, 1H and S1C). Osbp-YFP localized to endosomes, where we previously showed PI4KII functions (Burgess et al., 2012; Ma et al., 2020). Indeed, overexpressed Osbp-YFP decorated morphologically normal or enlarged endosomes in WT or *PI4KII*Δ salivary gland cells, respectively (Figures S1D and S1E). Thus, Osbp is a positive regulator of SG size, and its function in this process requires PI4KII.

**Figure 1.**
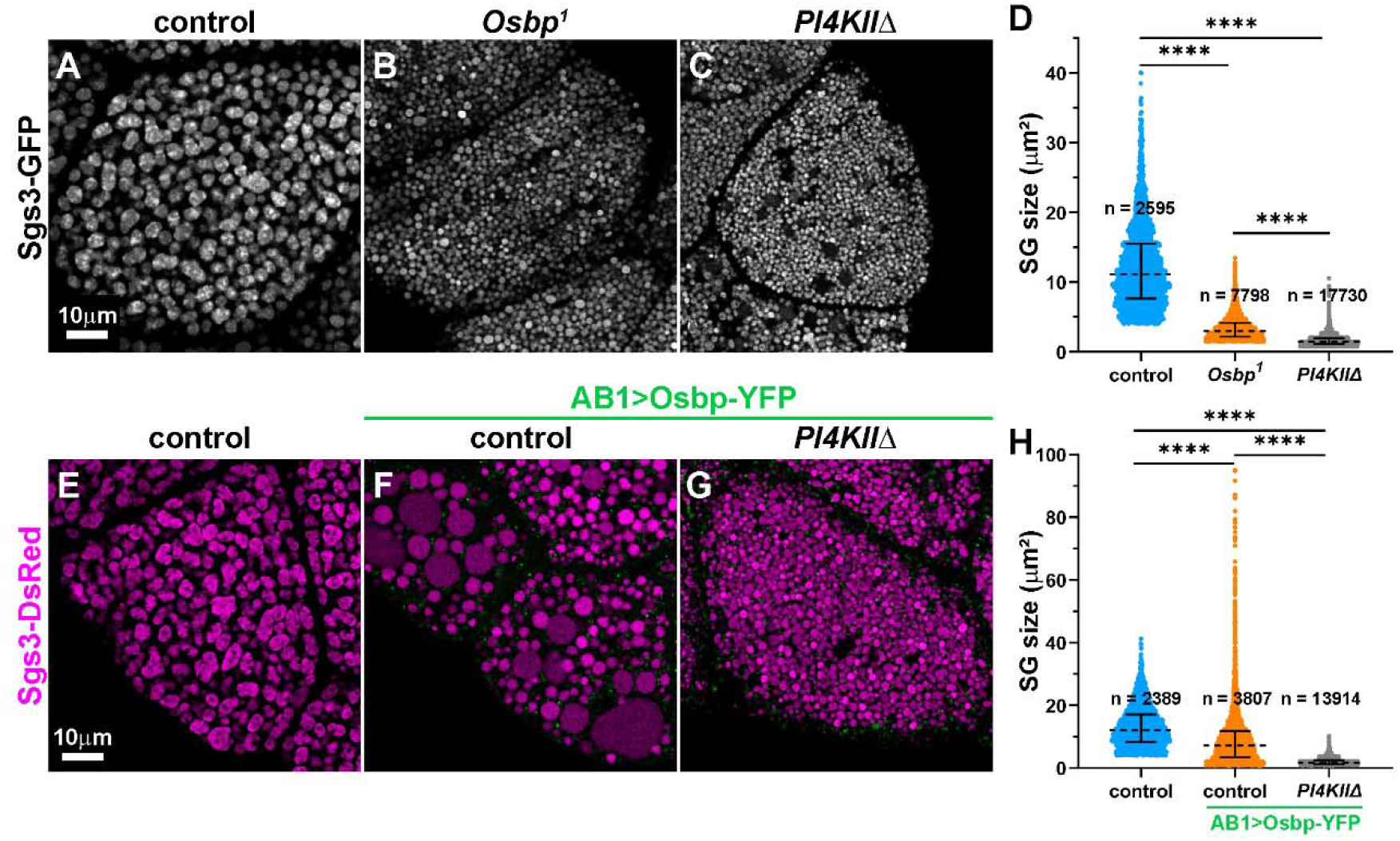
PI4KII is needed for Osbp to increase SG size. (A-C and E-G) Point-scanning confocal images of live late L3 salivary gland cells. Control (A), *Osbp^1^* (B), and *PI4KII*Δ (C) cells expressing Sgs3-GFP (grayscale). (D) Scatter plot showing the distribution of cross-sectional areas of SGs from A-C (see methods for details). n = total number of SG in images from three independent salivary glands (one salivary gland per independent experiment). Control (E) cells expressing Sgs3-DsRed (magenta). Control (F) and *PI4KII*Δ (G) cells expressing Sgs3-DsRed (magenta) and Osbp-YFP (green; driven by one copy of AB1-GAL4). (H) Scatter plot showing the distribution of cross-sectional areas of SGs from E-G (see methods). n = total number of SGs from three representative salivary glands (one salivary gland per independent experiment).

The PI4P phosphatase Sac1 and Vesicle-associated membrane protein-associated protein (Vap33) are positive regulators of Osbp function in *Drosophila* and mammals (Dong et al., 2016; Mao et al., 2019; Moustaqim-barrette et al., 2014). However, a temperature sensitive Sac1 mutant (*Sac1^ts^;* Del Bel et al., 2018; Griffiths et al., 2020) and *Vap33* RNAi salivary gland cells exhibited a large SG phenotype similar to Osbp overexpression (Figures S1F-S1K) rather than the small SG phenotype observed in *Osbp^1^*. Because loss of Sac1 or Vap33 can lead to an increase in PI4P levels (Forrest et al., 2013), we suspected that the large SG phenotype may be a result of elevated PI4P and that Osbp may also be involved in regulating PI4P levels and distribution.

To determine how Osbp affects PI4KII localization, we expressed GFP-PI4KII in control, *Osbp^1^*and Osbp overexpressing salivary gland cells. In control cells, GFP-PI4KII decorated membrane tubules that originated from endosomes, as previously described (Figure 2A; Burgess et al., 2012; Ma et al., 2020). In *Osbp^1^* cells, PI4KII-positive tubules were absent, and GFP-PI4KII accumulated on enlarged or irregularly shaped endosome-like structures (Figure 2B). In Osbp overexpressing cells, GFP-PI4KII decorated tubules emanating from enlarged endosomes (Figure 2C). To examine whether changes in PI4KII localization in *Osbp^1^* had consequences for PI4P distribution, we co-expressed the pan-PI4P biosensor mCh-2xP4M with Sgs3-GFP in salivary gland cells (Hammond et al., 2014; Ma et al., 2020). In control cells, mCh-2xP4M decorated the limiting membranes of SGs, as previously reported (Figures 2D; Ma et al., 2020). However, the level of PI4P on SG limiting membrane was significantly reduced in *Osbp^1^* cells, and most of the detectible PI4P accumulated on enlarged endosomes that contained a faint Sgs3-GFP signal (Figures 2E, 2G and 2H). In Osbp overexpressing cells, the level of PI4P on SG limiting membranes was significantly increased compared to controls (Figures 2F, 2G, and 2H). Because *Sac1^ts^* cells exhibited large SGs, similar to Osbp overexpressing cells, we examined whether a similar change would be observed in the level of SG-associated PI4P. Indeed, the level of PI4P on SG limiting membranes was also increased in *Sac1^ts^* cells (Figure S2). Therefore, PI4P on the limiting membranes of SGs positively correlates with SG size.

**Figure 2.**
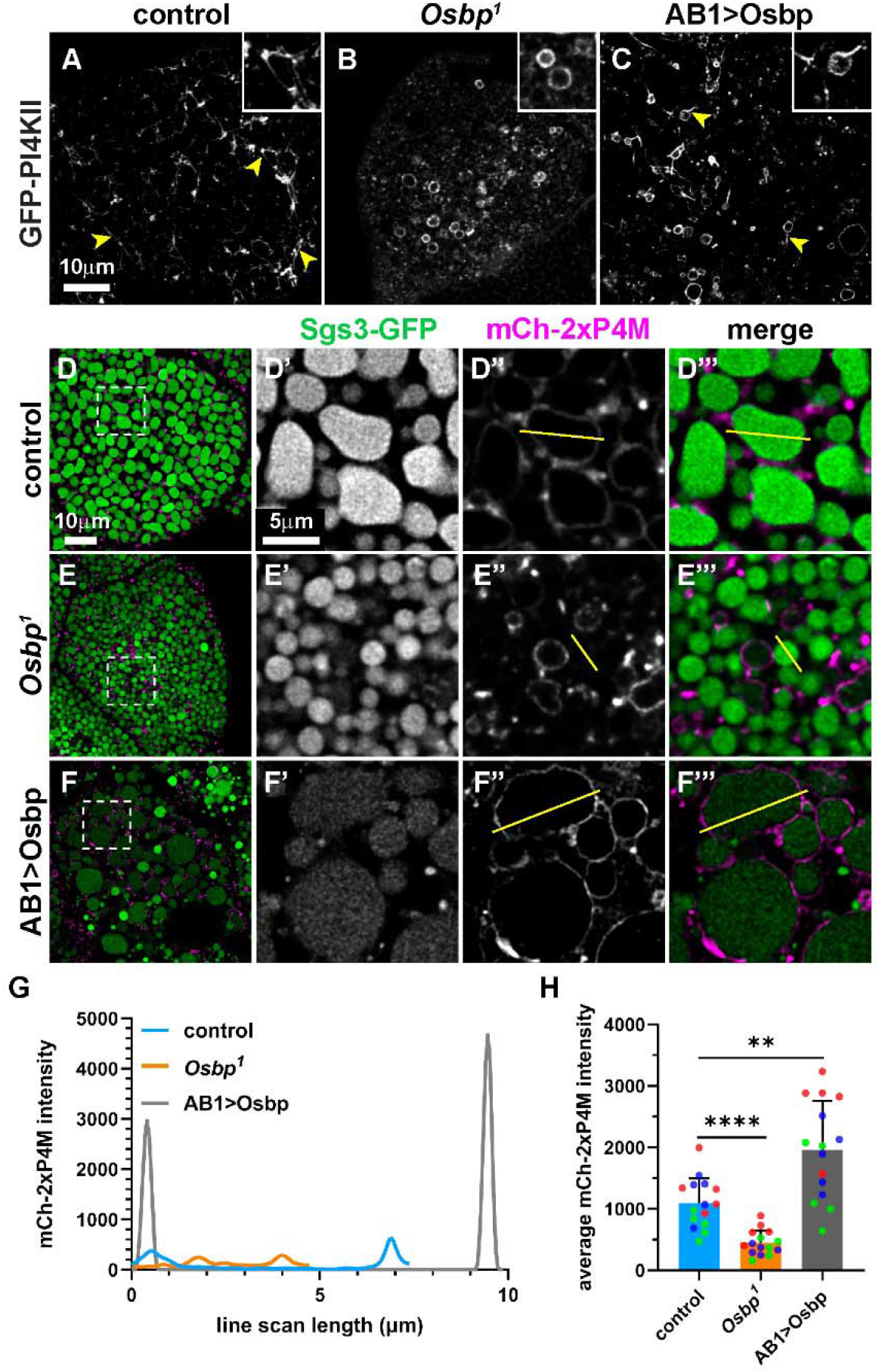
Osbp regulates PI4KII-dependent tubule formation and PI4P distribution. (A-F) Point-scanning confocal images of live late L3 salivary gland cells. (A-C) GFP-PI4KII localization in control (A), *Osbp^1^* (B), and Osbp overexpressing (C; driven by one copy of AB1-GAL4) cells. Yellow arrowheads indicate PI4KII tubules. (D-F) Cells expressing Sgs3-GFP (green) and mCh-2xP4M (magenta) from control (D), *Osbp^1^* (E), and Osbp overexpressing (F) salivary glands. Dashed boxes mark regions magnified 4.3-fold. Yellow lines depict representative line scans. (G) Line plot for mCh-2xP4M fluorescence intensity from representative line scans shown in D-F. (H) Bar graph showing average fluorescence intensity of mCh-2xP4M from line scans (see methods). *n* = 15 line scans from three representative salivary glands (one salivary gland per independent experiment, grouped by color).

Although Osbp overexpressing cells resulted in a subpopulation of very large SGs, these SGs appeared to have dimmer Sgs3-DsRed/GFP signals when compared to smaller SGs from the same cells (Figures 1F and 2F). The dimmer signal made us suspect that these aberrantly large SGs may result from an unregulated increase in size without a proportional increase in SG content, or that there might be excess fusion between endolysosomal and secretory compartments leading to degradation of SG content by lysosomal enzymes. Because both loss of Osbp and Osbp overexpression resulted in aberrantly large endosome-like structures marked by GFP-PI4KII (Figures 2B and 2C), we hypothesized that the enlarged endosomes in *Osbp^1^* and Osbp overexpressing cells were due to increased autophagy, and the large SGs may result from uncontrolled fusion between autolysosomes and SGs. To test this hypothesis, we stained control, *Osbp^1^* and Osbp overexpressing cells with anti-GABARAP antibodies, which detect Atg8a (LC3 in mammals) in *Drosophila* (Kim et al., 2015; Nandi et al., 2017). We observed an increase in GABARAP puncta in both *Osbp^1^* and Osbp overexpressing cells, suggesting an increase in autophagy initiation in these cells (Figures 3A-3C and 3G). Because formation of autolyososmal tubules is needed for autophagosome resolution (Munson et al., 2015; Rong et al., 2011; Yu et al., 2010), we suspected that loss of PI4KII-positive tubules (Figure 2B) may impact this process in *Osbp^1^*cells. To examine this, we stained control, *Osbp^1^* and Osbp overexpressing cells for Ref(2)P (*Drosophila* ortholog of the autophagy adaptor protein p62). Inhibition of autophagy or defects in autophagolysosome formation are correlated with accumulation of Ref(2)P that marks ubiquitinated proteins for autophagic or proteasomal degradation (Devorkin and Gorski, 2014; Nandi et al., 2017; Pircs et al., 2012). *Osbp^1^*cells had more Ref(2)P puncta that were also larger and brighter when compared to control and Osbp overexpressing cells, suggesting a defect in clearance of ubiquitinated proteins (Figures 3D-3G). In contrast, Osbp overexpressing cells showed no apparent increase in Ref(2)P levels, suggesting normal degradation (Figure 3F). Because Osbp overexpressing cells have an increase in autophagy initiation with normal degradation of Ref(2)P positive ubiquitinated proteins, the aberrantly large SGs observed in these cells may be a result of uncontrolled fusion between autophagolysosomes and SGs.

**Figure 3.**
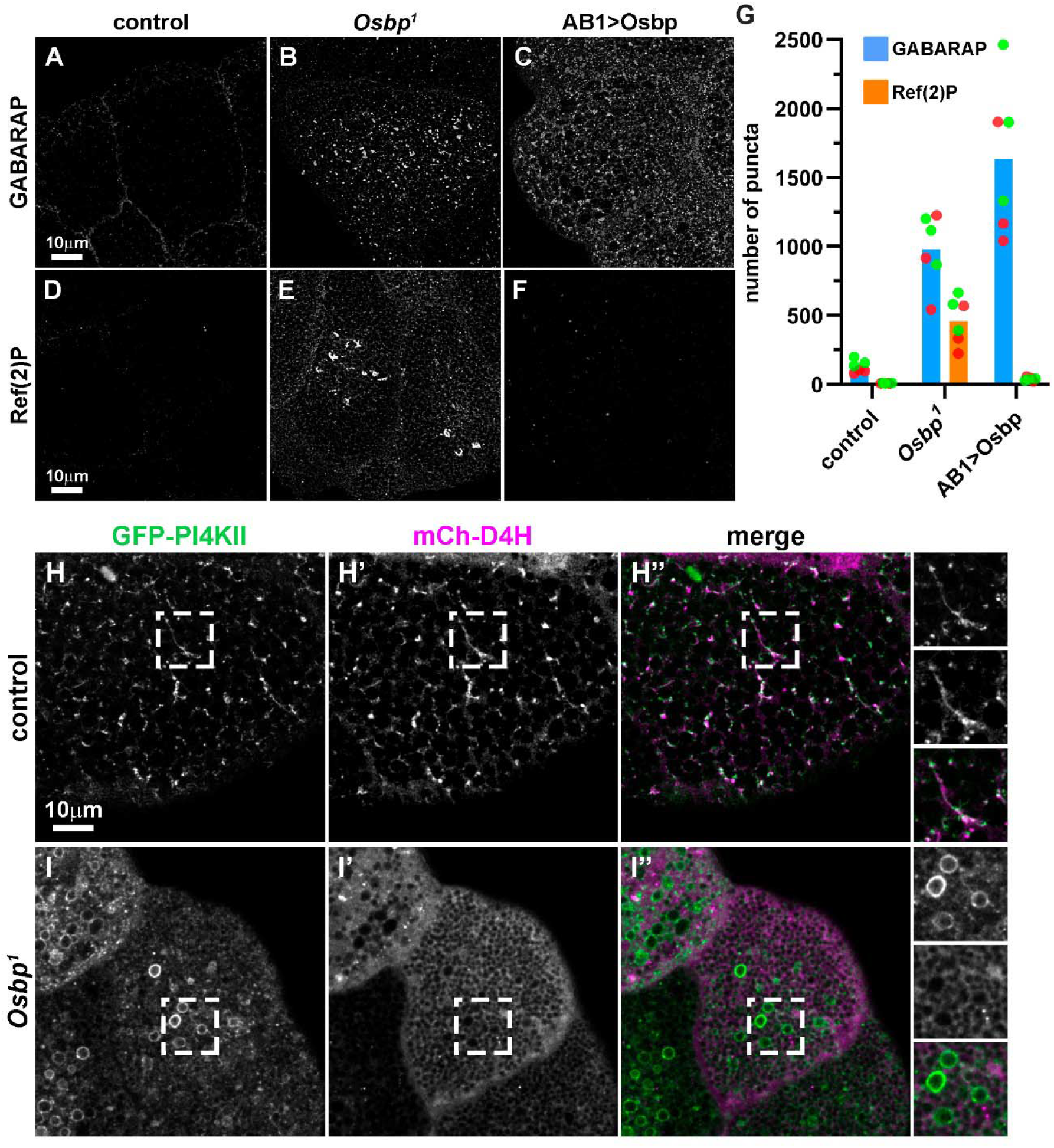
Osbp levels affect autophagy and autophagosome resolution. Point-scanning confocal images of (A-F) fixed or (H and I) live late L3 salivary gland cells. Immunostaining of GABARAP in control (A), *Osbp^1^*(B), and Osbp overexpressing (C) cells. Immunostaining of Ref(2)P in control (D), *Osbp^1^* (E), and Osbp overexpressing (F) cells. (G) Bar graph showing number of GABARAP and Ref(2)P positive puncta from A-F. Three fields of view were analyzed from two independent salivary glands (one salivary gland per independent experiment, grouped by color). Control (H) or *Osbp^1^* (I) cells expressing GFP-PI4KII (green) and mCh-D4H (magenta). Dashed boxes mark regions magnified 1.6-fold in insets.

Cholesterol levels can also affect autophagy and alter autophagic flux, as cholesterol depletion via methyl-β-cyclodextrin extraction or HMG-CoA (3-hydroxy-3-methyl-glutaryl-coenzyme A) reductase inhibition causes upregulation of autophagy and inhibition of autophagic flux in human and mouse fibroblasts (Cheng et al., 2006). To examine whether loss of Osbp had any effect on the distribution of membrane cholesterol on PI4KII-positive endosomes, we co-expressed the cytosolic cholesterol biosensor mCherry-D4H (mCh-D4H) with GFP-PI4KII in salivary gland cells (Burgess et al., 2012; Maekawa and Fairn, 2015). In control salivary gland cells, GFP-PI4KII and mCh-D4H colocalized to endosomes and membrane tubules that emanated from endosomes (Figures 3H), suggesting these endosomal membrane tubules are rich in cholesterol. In contrast, in *Osbp^1^*salivary gland cells, GFP-PI4KII decorated enlarged hollow endosomes that lacked mCh-D4H (Figures 3I), indicating these endosomes were relatively poor in cholesterol content as compared to controls. In addition, mCh-D4H distribution was more diffuse, indicating that cholesterol in the cytoplasmic leaflet of cellular membranes is reduced in *Osbp^1^*cells. Thus, Osbp mediated cholesterol transfer to endosomal membranes appears crucial for the formation of PI4KII-positive endosomal tubules and resolution of autophagy.

*Drosophila*, like other insects, is a sterol auxotroph and can obtain sterol only through dietary uptake (Clayton, 1964). Previous studies demonstrated that maintaining *Osbp^1^* flies on sterol-rich food containing 7-dehydrocholesterol (an intermediate metabolite of cholesterol) could suppress the spermatogenesis defects observed in *Osbp^1^* mutants (Ma et al., 2010, 2012). To evaluate whether sterol feeding could also suppress the defects in larval salivary gland cells, we raised control and *Osbp^1^*adult flies and larvae on regular or sterol-rich food containing 0.2 mg/g 7-dehydro-cholesterol. Sterol feeding partially suppressed the defects in SG maturation (Figures 4A-4D, and 4I) and PI4P distribution (Figures 4E-4H, and 4J) that were observed in *Osbp^1^*larvae fed a regular diet. Although a significant difference was observed in SG size between control and *Osbp^1^* raised on sterol-rich food, this was due to incomplete rescue of small SGs by sterol feeding (Figure 4I, box with dashed lines). Indeed, a majority of SGs from control and *Osbp^1^*larvae grown on sterol supplemented food had a similar size distribution (see median and interquartile range in Figure 4I), which explains why there was no significant difference when comparing the average size of SGs (Figure S3A). Moreover, line scan analysis revealed that sterol supplemented control and *Osbp^1^* larvae had similar PI4P levels on their SG membranes (Figure 4J). To determine whether sterol supplementation affected intracellular cholesterol distribution in salivary gland cells, we evaluated the distribution and intensity of mCh-D4H under different feeding conditions. mCh-D4H-positive structures increased in total intensity for both control and *Osbp^1^* cells under sterol-rich diet (Figures S3B-S3F). Interestingly, mCh-D4H-positive tubules were shorter in control cells under sterol supplement (Figures S3B and S3D), correlating with smaller granule size in sterol-fed control flies (Figures 4A, 4C and 4I). Thus, increasing cellular sterol content by feeding partially rescues defects in SG size and PI4P distribution observed in *Osbp^1^* salivary gland cells.

**Figure 4.**
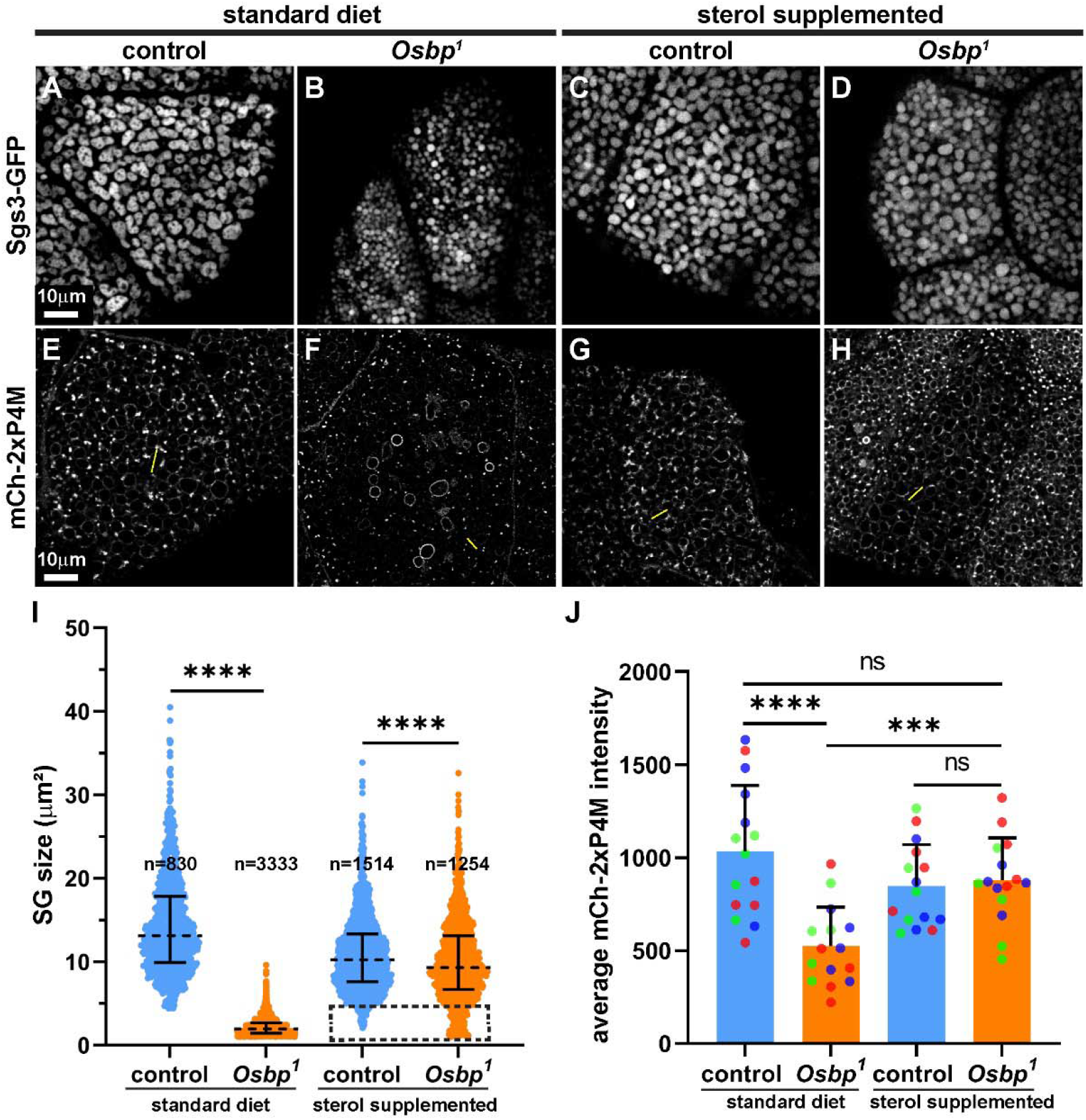
Sterol-rich diet partially suppresses *Osbp^1^* defect. (A-D) Spinning-disc and (E-H) point-scanning confocal images of live late L3 salivary gland cells. Cells expressing Sgs3-GFP (A-D; grayscale) or mCh-2xP4M (E-H; grayscale) from control and *Osbp^1^* larvae fed with a standard (left), or sterol supplemented (right) diet. Yellow lines depict representative line scans for measuring mCh-2xP4M intensity. (I) Scatter plot showing the distribution of cross-sectional areas of SGs from A-D (see methods). n = total number of SGs in images from three representative salivary glands (one salivary gland per independent experiment). (J) Bar graph showing average fluorescence intensity of SG-associated mCh-2xP4M from line scans in E-H (see methods). *n* = 15 line scans from three representative salivary glands (one salivary gland per independent experiment grouped by color).

## Discussion

In this study, we provide evidence that Osbp increases SG size in a PI4KII-dependent manner. Moreover, Osbp is needed for formation of PI4KII-positive endosomal tubules, a process we previously showed depends on PI4KII catalytic activity (Burgess et al., 2012; Ma et al., 2020). Thus, reciprocal regulation of Osbp and PI4KII is important to facilitate SG maturation. Studies on mammalian OSBP and OSBP related proteins (ORPs) have demonstrated that OSBP and ORPs mediate bi-directional transport of lipids between organelles at membrane contact sites (MCS) (Antonny et al., 2018; Mesmin and Antonny, 2016). For example, OSBP and ORP1L exchange cholesterol and PI4P across ER–*trans*-Golgi network and ER– endolysosome MCS, respectively (Dong et al., 2016; Goto et al., 2016; Levin-Konigsberg et al., 2019; Mesmin et al., 2013; Wijdeven et al., 2016; Zhao and Ridgway, 2017; Zhao et al., 2020). Humans have 12 ORP genes that are alternatively spliced into 16 protein products. Seven of these contain all the conserved domains found in Osbp: PIP-binding PH domain, ER-binding FFAT motif, and sterol-binding domain. In contrast, of the four *Drosophila* Osbp-related proteins, only Osbp has all three conserved domains (Ma et al., 2010). As a result, it is likely that *Drosophila* Osbp may have broader cellular functions as compared to its mammalian homologues.

Our observations that overexpressed Osbp localizes to endosomes and that *Osbp^1^* mutants exhibit endosomes marked with PI4KII but not cholesterol suggest that *Drosophila* Osbp acts at ER–endosome MCS. Loss of colocalization of the cholesterol sensor D4H and PI4KII on endosomes in *Osbp^1^*salivary gland cells further supports this idea. D4H labeling revealed that PI4KII-positive endosomal tubules are enriched in cholesterol, whereas loss of Osbp resulted in enlarged endosomes that lack both PI4KII tubules and D4H signal. Loss of Osbp did not abolish the recruitment of PI4KII to endosomes, and PI4KII remained active on the endosomes as demonstrated by accumulation of the PI4P sensor 2xP4M. Hence, endosomal cholesterol in *Osbp^1^* cells is apparently sufficient to recruit PI4KII and facilitate its production of PI4P (Barylko et al., 2009; Lu et al., 2012; Minogue et al., 2010; Waugh et al., 2006). However, tubulation of endosomal membranes failed to occur without Osbp facilitating additional transport of cholesterol to endosomes. Cholesterol could enable tubulation by helping to induce membrane curvature or increasing membrane elasticity (Döbereiner et al., 1993; Song and Waugh, 1993). Furthermore, cholesterol is needed for BAR domain proteins to tubulate liposomes *in vitro* (Daum et al., 2016). Previous studies have also demonstrated that endosomal membrane cholesterol is important for the recruitment of cytoplasmic phospholipase A2 and EHD1/Past1 to endosomes for tubulation and vesiculation (Cai et al., 2012). Therefore, it seems likely that reduced endosomal cholesterol content in *Osbp^1^* mutants affects endosomal flux via endosomal tubulation, thereby leading to the presence of enlarged endosomes. Having too much cholesterol in endosomal membranes may also affect tubulation, as we observed WT salivary gland cells under sterol-rich diet have shorter D4H-positive tubules when compared to cells under standard diet. This PI4KII and Osbp mediated retrograde trafficking may also be one of the many mechanisms that maintain intracellular cholesterol homeostasis. Golgi-associated retrograde protein (GARP) complex mediated retrograde transport may play a similar role, as loss of the GARP complex subunit Vps53 results in lysosomal dysfunction with abnormal distribution of sphingolipids and cholesterol similar to lipid storage diseases (Fröhlich et al., 2015; Wei et al., 2017).

We observed that loss of Osbp led to increased autophagy initiation, but clearance of ubiquitinated proteins was compromised. Similar observations have been made following knockdown of another OSBP family member, ORP1L, or treatment with an OSBP inhibitor, suggesting that proper regulation of cholesterol levels on endolysosomal membranes is important to resolve autophagosomes (Baba et al., 2019; Wijdeven et al., 2016). This defect was not observed upon Osbp overexpression, suggesting that Osbp levels may control resolution of autophagic cargoes or motility and fusion of lysosomes with other organelles. PI4P and cholesterol enriched membrane tubules could be important for the resolution of lysosomes and phagolysosomes since these tubules are absent in *Osbp^1^* mutants. This idea is supported by a previous study in mammalian macrophages demonstrating the importance of ORP1L in resolution of phagolysosomes in a PI4P dependent manner (Levin-Konigsberg et al., 2019). Furthermore, numerous studies in mammalian cells have reported that ORP1L forms a complex with Rab7 and Rab-interacting lysosomal protein (RILP), which induces recruitment of dynein-dynactin motor complexes that facilitate migration and fusion of LEs and lysosomes (Johansson et al., 2005, 2007; Levin-Konigsberg et al., 2019; Ma et al., 2018; Rocha et al., 2009; van der Kant et al., 2013; Vihervaara et al., 2011). Although the *Drosophila* RILP ortholog Rilpl is not well characterized, a previous study demonstrated that Rilpl associates with the lysosomal small GTPase Arl8 (Rosa-Ferreira et al., 2018). Moreover, expression of Rilpl in *Drosophila* S2 cells caused Arl8 to cluster near the perinuclear region (Rosa-Ferreira et al., 2018), consistent with previous observations in mammalian cells (Cantalupo et al., 2001; Jordens et al., 2001). Consistent with the idea that Osbp may share some properties of ORP1L and alter SG maturation by affecting motility of late endosomes and lysosomes, our previous observations indicate that PI4KII is also required for proper motility and distribution of endosomes (Burgess et al., 2012). Whether this is the case will require future study.

Raising *Osbp^1^* larvae on sterol-rich food suppressed SG maturation defects in salivary gland cells, similar to what was observed for sperm development in the adult testis (Ma et al., 2010). Dietary sterol is absorbed by the gut and redistributed to other tissues via lipoprotein particles (Blacklock and Ryan, 1994; Palm et al., 2012). Sterol then enters *Drosophila* cells through endocytosis of these particles. Lipids are hydrolyzed in lysosomes before reaching equilibrium in the ER, where they can be stored in lipid droplets or redistributed to other organelles. Thus, it is possible that the partial suppression of the *Osbp^1^* phenotype by sterol feeding is due to elevating sterol levels in membranes of multiple intracellular organelles. Moreover, SGs in mammalian cells are enriched in cholesterol, and multiple studies have also shown that cholesterol levels can impact SG size and maturation in mammalian secretory cells (Bogan et al., 2012; Gondré-Lewis et al., 2006; Hussain et al., 2018; Orci et al., 1981; Sun et al., 2013). Interestingly, cholesterol loading restores insulin granule maturation defects in rat insulinoma cells with reduced ABCA1 and ABCG1 levels, but a similar effect is not observed in cells with reduced levels of OSBP (Hussain et al., 2018). Nevertheless, our results provide further evidence that fine tuning of sterol levels is crucial for SG maturation.

Mesmin and colleagues described a mechanism of mutual dependence between OSBP and PI4Ks in immortalized human retinal pigment epithelial cells (Mesmin et al., 2017). Golgi localized PI4KIIIβ provides PI4P to enable OSBP-mediated cholesterol transport from the ER to the TGN. Palmitoylated PI4KIIα is then recruited the newly cholesterol enriched site to replenish PI4P for additional transport. In this study, we demonstrate for the first time that mutual dependence between PI4KII and Osbp for SG maturation occurs *in vivo* in *Drosophila* larval salivary glands. However, PI4KII and Osbp do not directly mediate PI4P production on SGs in salivary gland cells (Ma et al., 2020; and this work). Instead, PI4KII produces PI4P on endosomes and Osbp exchanges PI4P to enrich endosomal membranes with sterol. Transfer of sterol into endosomal membranes is crucial for the formation of sterol- and PI4P-rich tubules. These sterol- and PI4P-rich endosomal tubules then, directly or indirectly, deliver PI4P to SG membranes to mediate maturation. Loss of either Osbp or PI4KII abolishes formation of these tubules, resulting in immature SGs. Moreover, Golgi localized PI4KIIIβ is not involved in this process because loss of PI4KIIIβ (Fwd in *Drosophila*) does not affect SG maturation (Burgess et al., 2012). Our study provides further mechanistic insights into PI4P regulation of SG biogenesis *in vivo* using *Drosophila* larval salivary glands as a model. Defects in granule maturation could result in the release of secretory cargoes with subpar biological activities, leading to impaired homeostasis and disease states. For example, elevated proinsulin secretion is associated with both type 1 and type 2 diabetes, and impairment in von-Willebrand factor maturation can cause bleeding disorders (Huang et al., 2021; Starke et al., 2013; Steenkamp et al., 2017). Therefore, understanding how membrane lipids regulate SG biogenesis may provide key insights in the development of novel therapeutics to treat diseases associated with defects in regulated secretion.

## Materials and methods

### Fly Genetics

Flies were cultured on standard cornmeal molasses agar at 25°C (Ashburner, 1989). Fly stocks were acquired from Bloomington Drosophila Stock Center (BDSC) unless otherwise stated. Stocks used in this study include *P{GawB}AB1*; *P{w^+^, Sgs3-GFP}* (Biyasheva et al., 2001); *P{w^+^, UAS-Osbp-EYFP}*; *P{w^+^, UAS-Osbp.m}3*; *P{TRiP.JF02621}attP2* (Vap33 RNAi); *P{w^+^, CG14671}*, *P{w^+^,* α*Tub84B-mCh-2xP4M}* (Ma et al., 2020); *P{w^+^,* α*Tub84B-mCh-D4H}* (this study); *P{w^+^,* α*Tub84B-GFP-PI4KII}* (Burgess et al., 2012); *P{neoFRT}82B, Df(3R)730* (*PI4KII*Δ; Burgess et al., 2012); *w^1118^;;Osbp^1^*(Ma et al., 2010; a gift from X. Huang, Chinese Academy of Sciences, Beijing, China); *P{w^+^, Sgs3-DsRed}* (Costantino et al., 2008; a gift from A. Andres, University of Nevada, Las Vegas, NV, USA); *w^1118^;;Sac1^ts^*(Del Bel et al., 2018). UAS lines were expressed in salivary gland cells under control of the AB1-GAL4 driver (*P{GawB}AB1*). Controls used in this study were *w^1118^*flies carrying specified fluorescent transgenes.

### Molecular Biology

mCh-D4H sequences were PCR-amplified from a mammalian vector containing mCh-D4H (Maekawa and Fairn, 2015; a gift from G. Fairn, St. Michael’s Hospital, Toronto, ON, Canada) and subcloned into a modified pCaSpeR4 vector containing the αTub84B promoter (Marois et al., 2006) using In-Fusion cloning kit (Takara Bio, Japan). Forward primer sequence was 5’-CTAGAGGATCCCCGGGTACCATGGTGAGCAAGGGCGAG-3’ and reverse primer sequence was 5’-TCGAGGGGGGGCCCGGTACCTTAATTGTAAGTAATACTAGATCCAGGGTATAAAGTT GTTC-3’. Primers were synthesized by Thermo Fisher Scientific (USA).

### Generation of Transgenic Flies

mCh-D4H lines were generated by P-element transformation, wherein the plasmid containing αTub84B-mCh-D4H was injected into *w^1118^* embryos by BestGene Inc. (Chino Hills, CA, USA)

### Microscopy

For live imaging, salivary glands were dissected from late L3 larvae in 50 µL Drosophila Ringer’s solution (10 mM Tris, 182 mM KCl, 46 mM NaCl, 3mM CaCl_2_·2H2O, pH 7.2) using fine forceps (Fine Science Tools #11251-20, Canada) . Dissected salivary glands were transferred to 8 µL Drosophila Ringer’s on a microscope slide with self adhesive reinforcement labels (Avery #32203, USA) as spacers and sealed using nail polish (Ma and Brill, 2021b). Samples were imaged using a Quorum spinning disc confocal coupled with an Olympus IX81 microscope (Quorum Technologies Inc., Canada), or a Leica SP8 point scanning confocal microscope (Leica, Germany). Images were acquired from the spinning disc confocal using a 60X oil objective (NA 1.4) and Volocity 6.3 (Quorum Technologies Inc., Canada) software. Serial optical sections were acquired at an interval of 0.3 µm for a total of 20-30µm. Images were acquired from Leica SP8 Lightning confocal using a 40X oil objective (NA 1.3) and LAS-X software (Leica, Germany). Leica SP8 images were processed with LIGHTNING using adaptive strategy from the LAS-X software. All images from the same experiment were adjusted for brightness and contrast in an identical manner using Adobe Photoshop CC (Adobe, USA).

### Immunofluorescence Microscopy

For immunofluorescence, salivary glands were dissected from late L3 larvae in 50 µL phosphate buffered saline (PBS, pH 7.4). Dissected glands were fixed with 4% paraformaldehyde PBS for 20 min at room temperature and permeabilized in PBST (PBS + 0.1% Triton X-100). Primary antibody incubation was performed overnight at 4°C in PBST with 5% normal goat serum (Thermo Fisher, USA). Secondary antibodies were diluted in PBST and incubated for 1 hr at room temperature. Samples were mounted in 8 µL ProLong Glass (Thermo Fisher, USA) using self adhesive reinforcement labels (Avery #32203, USA) as spacers. Mounted samples were cured overnight at room temperature before sealing with nail polish. Primary antibodies were rabbit anti-Ref(2)P (1:1000; a gift from T. E. Rusten, University of Oslo, Norway), rabbit anti-GABARAP (1:200; Abcam ab109364, UK). Secondary antibodies conjugated to Alexa Fluor 488 were used at 1:500 (Thermo Fisher, USA).

### Sterol Feeding Assay

Flies expressing Sgs3-GFP, mCh-2xP4M, or mCh-D4H were raised as described above with or without additional sterol (0.2 mg/g 7-dehydrocholesterol in ethanol or equal volume ethanol). Salivary glands of late L3 larvae were dissected and examined live on a spinning-disc or point scanning confocal microscope, as described above.

### Image Analysis

Cross-sectional areas of SGs were quantified using Volocity 6.3 as previously described (Ma and Brill, 2021b; Ma et al., 2020). Images were first deconvolved using the iterative restoration function with a confidence level of 99% and a maximum of 30 iterations. The find object function was used to identify and measure the surface area of SGs. Objects with a cross-sectional area less than 4 µm^2^ or 0.8 µm^2^ were filtered out for control or *Osbp^1^* salivary glands, respectively. Analyses were performed on two to three z-slices from a single acquisition (∼3-4 cells per field of view) from three individual salivary glands. Quantification for sterol feeding assay was limited to one cell per field of view because not all neighboring cells were rescued.

Line scan analysis was performed using Fiji-Image J software by drawing a line across SG to measure the intensity in the mCh-2xP4M channel. Intensities over distance were plotted as a line plot. The two local maxima of mCh-2xP4M intensities along a single line were averaged and plotted as a data point for the average fluorescence intensity bar graph.

Number of puncta were counted using the find object function in Volocity 6.3 by applying global and local thresholding.

### Statistical Analysis

Raw data from ImageJ and Volocity 6.3 were imported into Microsoft Excel 365 for data processing. Final numbers were imported to GraphPad Prism 9 (GraphPad Software, USA) to be plotted and analyzed. Student’s t-tests (two-tailed, unequal variance) were performed to test for statistical significance. *p < 0.5; **p < 0.01; ***p < 0.001; ****p < 0.0001.

Scatter plots were generated using GraphPad Prism 9. The center line represents median, whereas the error bar represents interquartile range (25^th^ and 75^th^ percentile).

---

**Supplemental Figure 1.**
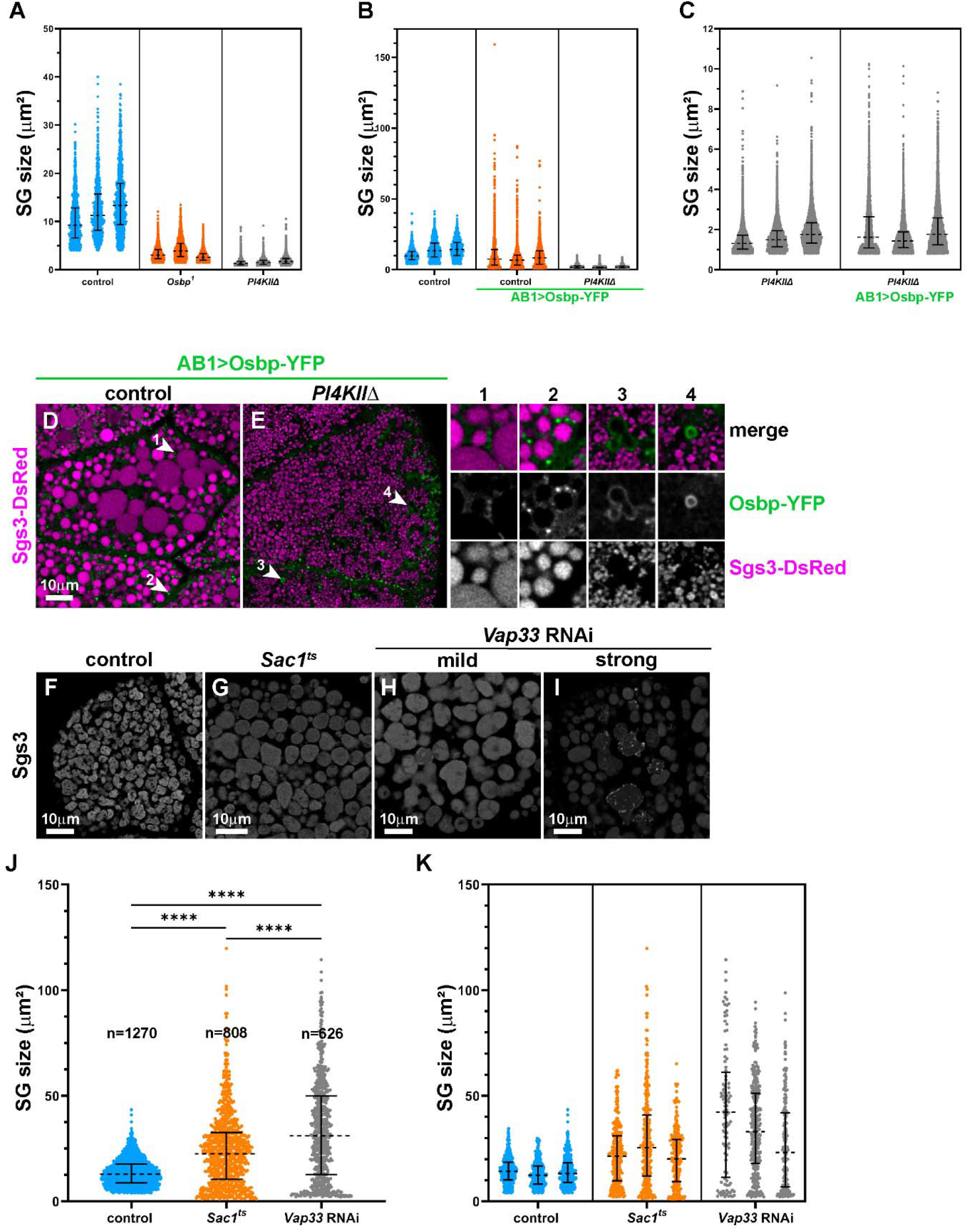
Osbp localization and SG phenotypes in *Sac1^ts^* and *Vap33* RNAi cells. (A-B) Scatter plots showing the distribution of cross-sectional areas of SGs from each independent salivary gland in Fig. 1 D and H (see methods). (C-H) Point-scanning confocal images of live late L3 salivary gland cells. Control (C) and *PI4KII*Δ (D) cells expressing Sgs3-DsRed (magenta) and Osbp-YFP (green; driven by one copy of AB1-GAL4). White arrowheads 1-3 mark regions magnified ∼2.4 fold in insets. Cells expressing fluorescently-tagged Sgs3 (grayscale) in control (E), *Sac1^ts^* (F), and *Vap33* RNAi (G-H) expressing cells. (I) Scatter plot showing the distribution of cross-sectional areas of SGs from E-H (see methods). n = total number of SG in images from three representative salivary glands (one salivary gland per independent experiment). (J) Scatter plots showing the distribution of cross-sectional areas of SGs from each independent salivary gland in I. (see methods).

**Supplemental Figure 2.**
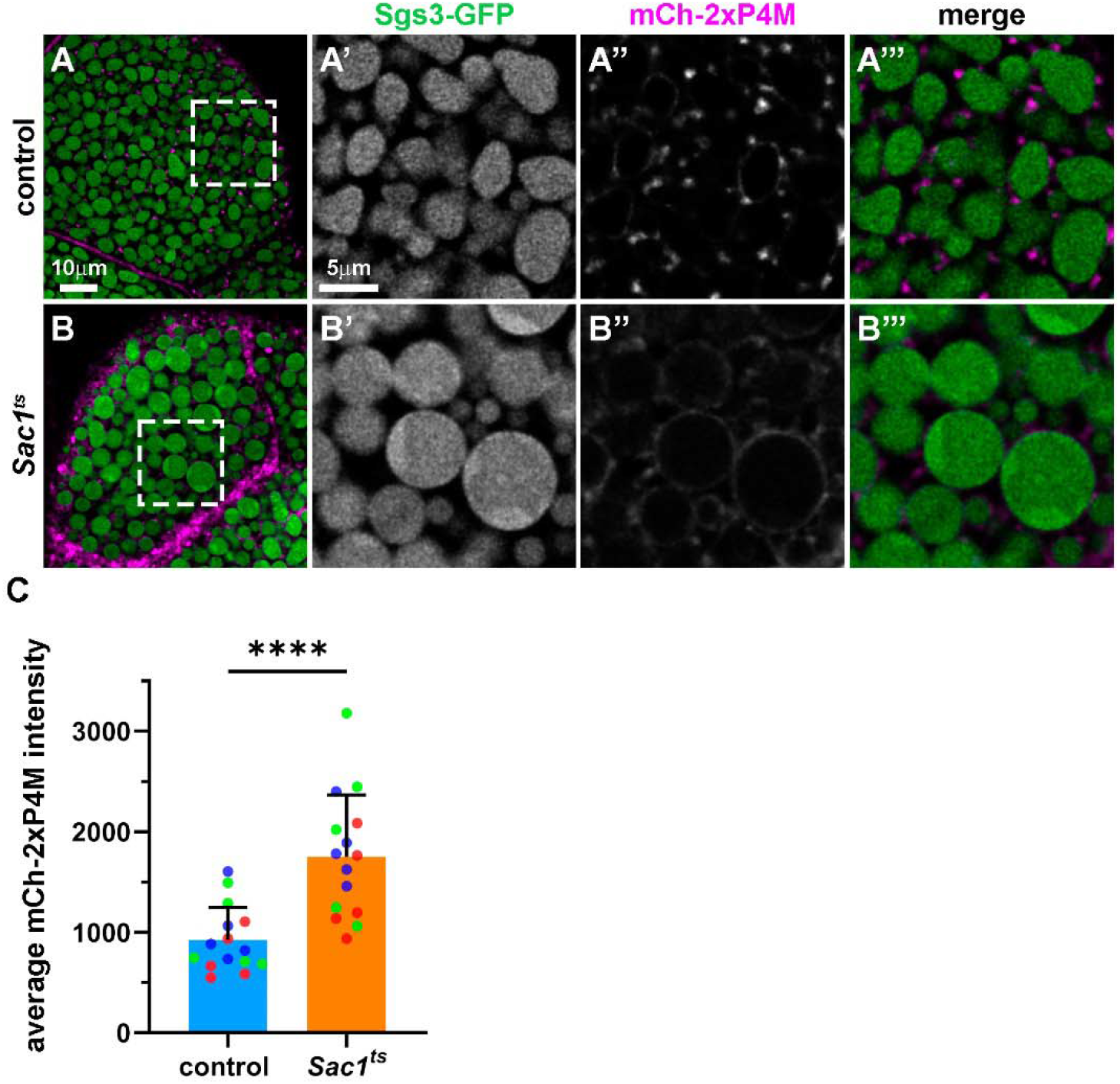
PI4P level on SG membrane is elevated in *Sac1^ts^* cells. Point-scanning confocal images of live late L3 salivary gland cells expressing Sgs3-GFP (green) and mCh-2xP4M (magenta) from control (A) and *Sac1^ts^* (B) larvae. (C) Bar graph showing the average peak fluorescence intensity of mCh-2xP4M from line scans (see methods). *n* = 15 line scans from three representative salivary glands (one salivary gland per independent experiment grouped by color).

**Supplemental Figure 3.**
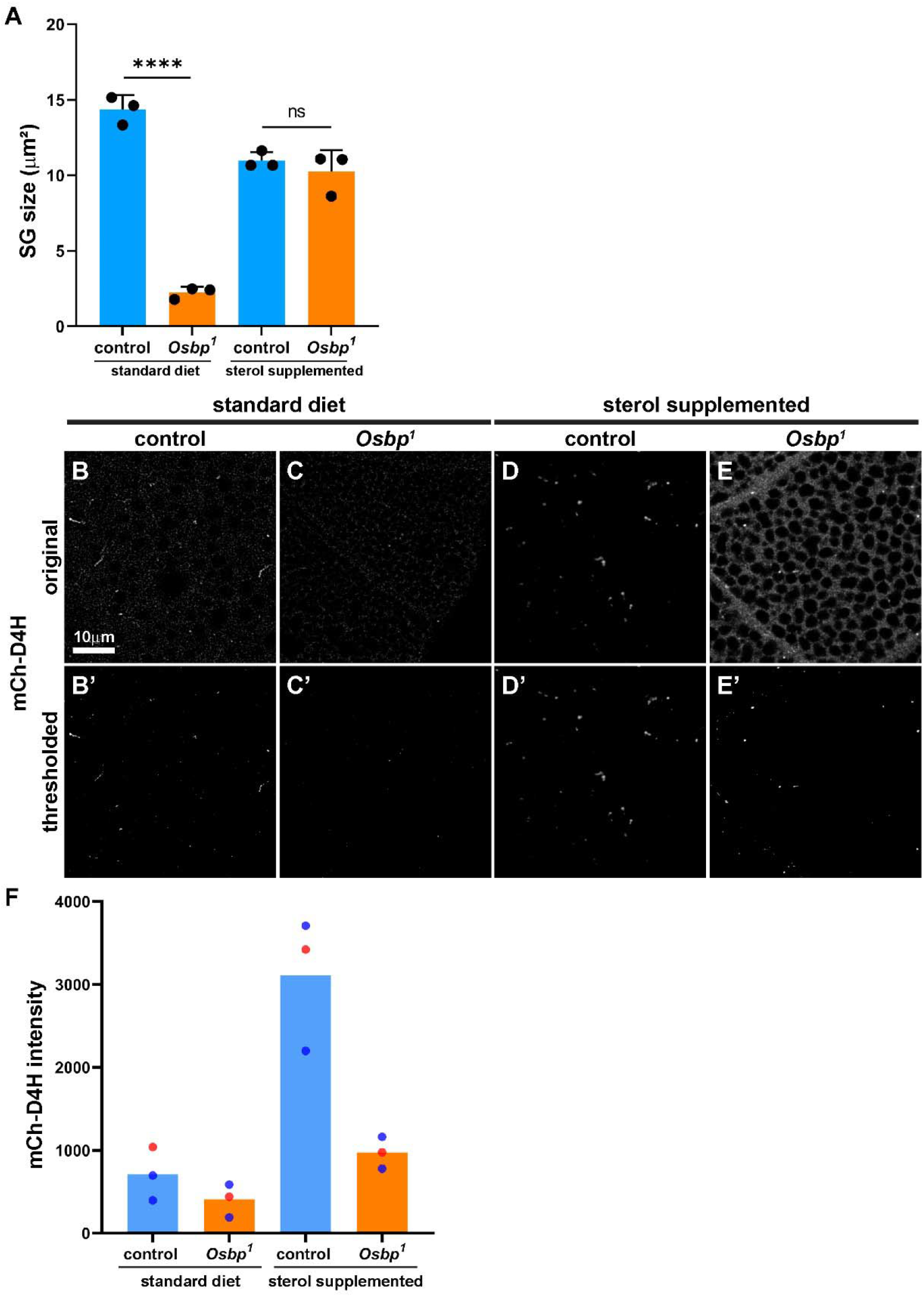
mCh-D4H levels increases under sterol-rich diet. (A) Bar graph showing average SG cross-sectional area from Fig. 4 A-D. Black circles represent averages from independent salivary glands. (B-E) Point-scanning confocal images of live late L3 salivary gland cells expressing mCh-D4H (grayscale) from control and *Osbp^1^*larvae under standard (left) or sterol supplemented (right) diet. (B’-E’) Images from B-E thresholded for bright objects above background. (F) Bar graphs depicting integrative intensity of thresholded objects. Each circle represents one field of view (3-4 cells per field of view) from two representative salivary glands (one salivary gland per independent experiment, grouped by color).

## Acknowledgements

We thank J. Brumell, U. Tepass, G. Fairn, H. Krämer, P. Kim, F. Maxfield and Brill lab members for helpful discussions; X. Huang, A. Andres and the Bloomington Drosophila Stock Center for fly stocks; G. Fairn for D4H plasmids; T. Rusten for anti-Ref(2)P antibodies; P. Paroutis and K. Lau for assistance with microscopy; Quorum Technologies Inc., Canada for providing remote access to Volocity imaging analysis software; FlyBase for providing a centralized resource for the *Drosophila* community; and BioRender.com for graphical abstract.

## Disclosure of Interest

The authors report no conflict of interest.

## Funding

This research was supported by the following funding sources: SickKids Restracomp Studentship, CIHR Strategic Training Fellowship #TGF-53877, University of Toronto Open Fellowship, and NSERC PGS-D Scholarship (to C-I. J. Ma) and CIHR Grants #PJT-162165 and #MOP-119483 (to J. A. Brill).

## Notes

### Competing Interest Statement

The authors have declared no competing interest.

